# MD simulations of Human Sigma1 Receptor Trimer Uncovers Cholesterol Dependent Stabilization and Ligand Specific Dynamics

**DOI:** 10.64898/2026.03.05.709821

**Authors:** Vittoria Nanna, Costanza Paternoster, Alessio Bartocci, Dritan Siliqi, Domenico Alberga, Carmen Abate, Gianluca Lattanzi, Giuseppe Felice Mangiatordi

## Abstract

The sigma-1 receptor (S1R) is an endoplasmic reticulum transmembrane protein implicated in a wide range of physiological and pathological processes, including neurodegeneration, cancer, and pain modulation. Although X-ray crystallography has revealed S1R as a trimeric assembly with a distinctive triangular architecture, the dynamic behavior of this oligomeric state and its modulation by ligands and membrane composition remain poorly understood. In particular, agonists and antagonists have been experimentally proved to differentially regulate S1R oligomerization although the underlying molecular mechanisms are still obscure. Here, we present the first atomistic molecular dynamics study of trimeric S1R embedded in a physiologically relevant lipid environment. Using a total of 12 µs of simulation time, we investigate the impact of membrane composition, with a specific focus on cholesterol, as well as the conformational response of S1R to pharmacologically distinct ligands: the agonist (+)-pentazocine and the antagonist haloperidol. Our simulations reveal how ligands can alter S1R interprotomer interaction through a mechanism involving the β6-strand of the protein and in particular W136, data that correlate with experimentally observed differences in S1R oligomerization. These findings provide new molecular-level insights into S1R regulation and establish a framework for rationalizing the distinct functional outcomes induced by agonists and antagonists.

## Introduction

The sigma-1 receptor (S1R) is a unique transmembrane protein located predominantly in the endoplasmic reticulum (ER) membrane, particularly at the critical mitochondria-associated membrane (MAM) junction where it orchestrates diverse cellular processes from calcium signaling to protein homeostasis.^1^ Its central role in neuroprotection, cancer progression, and pain modulation has positioned S1R as a compelling pharmacological target. In particular, S1R has been implicated in the progression of neurodegenerative diseases, including Alzheimer’s and Parkinson’s.^2^

To understand the diverse biological functions of S1R, it is essential to examine its peculiar molecular architecture. Structurally, S1R has been characterized *via* X-ray crystallography as a homotrimer composed of three 24-kDa subunits arranged in a triangular shape; each subunit presents a single N-terminal transmembrane helix positioned at the vertices of the triangle, and a cytoplasmic cupin-like β-barrel that contains the ligand-binding pocket (**Figure 1A**).^3^ This triangular assembly creates a flat, membrane-embedded platform with three symmetrically positioned binding sites. While this crystallographic snapshot provides crucial insight into overall fold of the receptor, it also raises new questions about the protein’s dynamic behavior. Experimental evidence suggests that S1R assembly is not rigid and static but rather a dynamic structure whose oligomerization state responds to ligand binding.^4–6^ In particular, agonists, such as (+)-pentazocine (hereinafter referred to as PnT), are thought to promote the dissociation of monomers from the assembly, while antagonists, like haloperidol (Hal), tend to stabilize the trimer and promote high-order oligomer formation.

**Figure 1.**
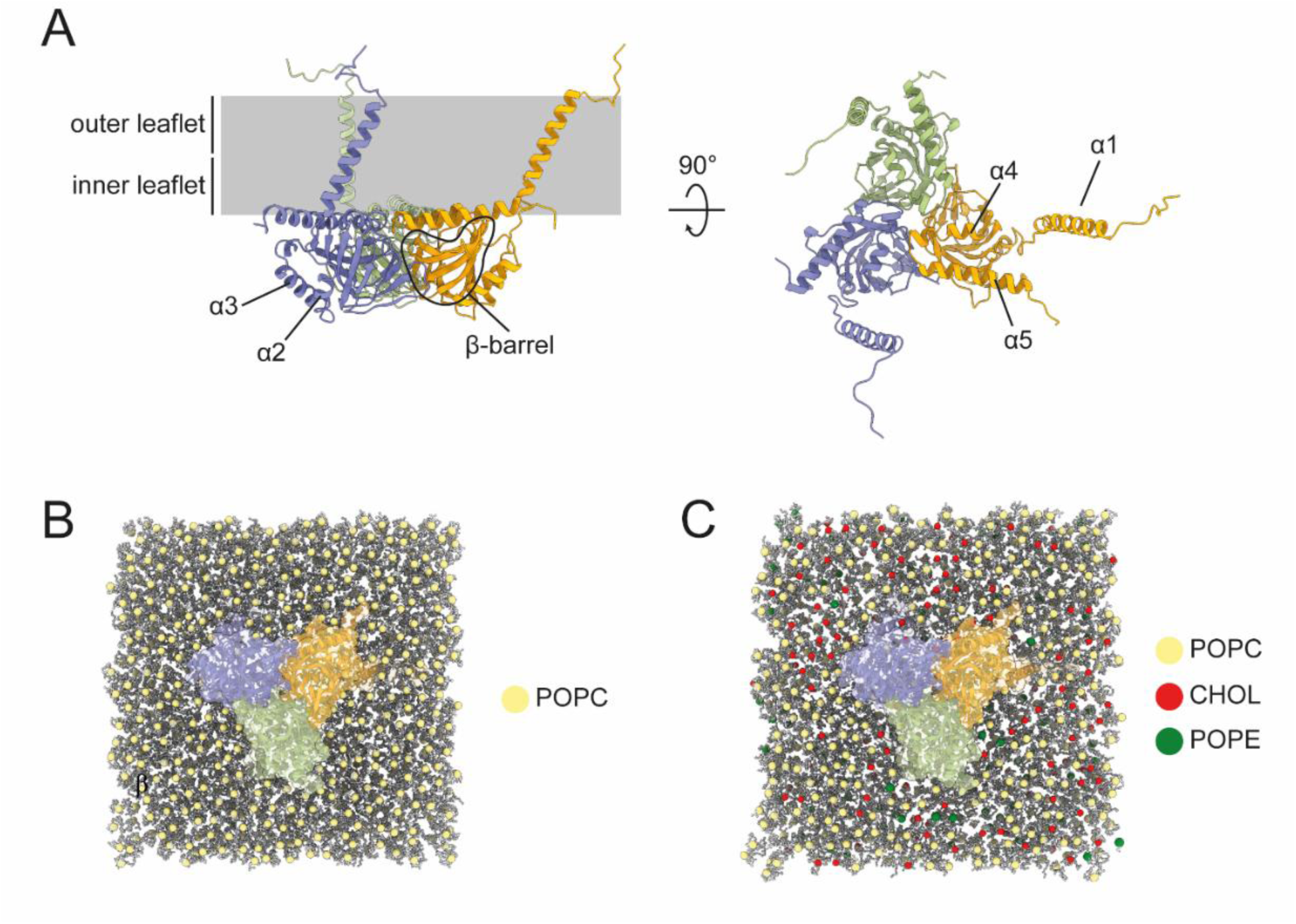
Structural features of S1R. **(A)** Front and top view of the human S1R structure (PDB code: 6DJZ^9^). The three monomers (A, B and C - colored in green, orange and purple, respectively) are arranged in a triangular shape. Secondary structure elements are labeled. **(B, C)** Representation of S1R embedding in the membrane in simulated systems. The pale-yellow dots represent the phosphate group of POPC, the red dots represent the oxygen group of cholesterol, and the green dots represent the phosphate group of POPE. Note that CHOL and POPE are equally distributed in the membrane in the initial setup.

Molecular dynamics (MD) simulations have been instrumental in exploring ligand binding of S1R. However, previous studies have largely focused almost exclusively on monomeric forms of the receptor embedded in simplified membrane models, typically composed of phosphatidylcholine (POPC) alone.^7,8^ While ligand-receptor interactions^9–11^ and ligand entry^12–14^ have been investigated in detail, the influence of binders on the receptor oligomeric state has not been sufficiently addressed.

In addition, experimental evidence points to a key role for the membrane environment in modulating S1R function. Cholesterol has been shown to directly interact with S1R and influence its behavior in the ER membrane.^15^ More recently, S1R has been implicated in shaping the ER membrane, with its trimeric assembly appearing essential to this function.^16^ Despite these insights, no atomistic simulations have yet addressed the dynamics of trimeric S1R. In this study, we fill this gap by performing extensive unbiased MD simulations (12 µs of trajectory data) to investigate: (i) how membrane composition influences the S1R behavior in its trimeric form, and (ii) the relationship between ligand activity (agonist *versus* antagonist) and the conformational response of S1R. To the best of our knowledge, this work represents the first atomistic MD investigation of human S1R trimer embedded in a physiologically relevant lipid environment, providing unprecedented insights into the role of cholesterol in receptor stability and into the distinct conformational responses induced by two pharmacologically different ligands, the agonist (+)-pentazocine^17^ and the antagonist haloperidol.^17^ Ultimately, the data presented here lay the groundwork for molecular-level hypotheses aimed at rationalizing the experimentally observed, yet mechanistically unresolved, differential effects of agonists and antagonists on S1R oligomerization.

## Materials & Methods

### Model system preparation

The structures used for the MD simulations were obtained from the Protein Data Bank (PDB), corresponding to S1R bound to haloperidol (PDB ID: 6DJZ) and to S1R bound to (+)-pentazocine (PDB ID: 6DK1).^9^ Henceforth, we refer to these complexes as S1R–Hal (S1R trimer bound to haloperidol) and S1R–PnT (S1R trimer bound to (+)-pentazocine). Modeller^18^ was employed to reconstruct the missing loops while the Protein Preparation Wizard (Schrödinger Suite, 2024-1)^19^ was used to reconstruct missing side chains and define the protonation state of titrable residues at physiological pH. Specifically, E172 was set in its charged state to interact with the ligands, whereas D126 was protonated to form the hydrogen bond with E172, as observed in structural studies.^3^

The S1R homo-trimer was embedded into a membrane bilayer composed of (i) exclusively phosphatidylcholine (POPC) and (ii) POPC (56%), phosphatidylethanolamine (POPE) (22%), and cholesterol (CHOL) (22%) reflecting the lipid composition of the MAM, which is enriched in cholesterol compared to the rest of the endoplasmic reticulum (**Figures 1B** and **1C**). This ratio was chosen based on the reported ER lipidic composition^20^ and the estimated cholesterol enrichment in the MAM.^21,22^ The systems were built using the Orientations of Proteins in Membranes (OPM) database for proper alignment along the x-y plane^23^ and constructed with the CHARMM-GUI web server.^24,25^ The resulting membranes formed asymmetric bilayers, with fewer lipids in the inner leaflet – directly contacting the protein core – than in the outer leaflet (**Supplementary Table S1**). This asymmetry was unavoidable and arose from the imposed orientation constraints. Specifically, maintaining the correct receptor orientation requires the insertion of the hydrophobic regions of helices α4 and α5, which lie parallel to the membrane plane. Each system was solvated with TIP3P water molecules, and Na^+^ and Cl^-^ ions were added to neutralize the net protein charge and to achieve a physiological salt concentration of 0.15 M. The final systems contained approximately 370,000 atoms. Detailed information on simulation box dimensions is provided in **Supplementary Table 2**. All simulations were performed using GROMACS version 2023 with the CHARMM36m force field for proteins, lipids, and solvent, and the CHARMM General Force Field (CGenFF) for ligand parameterization.^26–28^

### Simulation Protocol

The prepared systems were subjected to a steepest-descend minimization followed by a two-step equilibration protocol: 375 ps of simulation in the NVT ensemble (T=310 K, Berendsen thermostat^29^) with a 1 fs timestep, followed by 1.5 ns in the NPT ensemble (T=310 K and P=1 atm, Berendsen thermostat and barostat^29^) with a 2 fs timestep. Harmonic restraints were applied to protein and ligand heavy atoms, as well as on the z-coordinate of lipid phosphate atoms, in order to preserve bilayer thickness and to the acyl chain dihedrals. These restraints were then gradually released during throughout the equilibration phase. Initial velocities were randomly generated from a Maxwell-Boltzmann distribution at 310 K. Three independent replicates were simulated for each system (twelve replicates in total). Replicate 1 was initiated from the final snapshot of the restrained equilibration, while replicates 2 and 3 were obtained by extracting frames at 100 and 110 ns, respectively, from replicate 1, and assigning new velocities sampled from the Maxwell-Boltzmann distribution. The simulations were performed in the NPT ensemble, with temperature controlled at 310 K using a velocity rescaling thermostat (v-rescale) with a stochastic term^30^, with a coupling constant of τ_T_ = 1 ps, and pressure controlled at 1 bar using a c-rescale barostat^31^ with a coupling constant of τ_p_ = 5 ps and semi-isotropic coupling. To prevent membrane instabilities observed with default settings, which could be particularly pronounced in our system due to leaflet asymmetry, neighbor list and pressure-coupling parameters were carefully optimized following Kim et al.^32^; the full set of adopted parameters is reported in **Supplementary Table 3**. Long-range electrostatic interactions were computed with the particle-mesh Ewald method^33,34^ and non-bonded interactions were cut off at 1.2 nm. The LINCS algorithm was used to constrain the bonds containing hydrogen atoms. A time step of 2 fs was adopted, along with periodic boundary conditions. Each replicate of S1R was simulated for 1000 ns. For all considered systems, S1R equilibration required less than 200 ns and thus the first 200 ns were removed from the analysis.

### Analyses of the trajectories

The analyses were performed by sampling the trajectories every 500 ps when not explicitly specified. GROMACS utilities were used to manipulate the trajectories and compute the root mean square deviation RMSD (*gmx rms*). The *LipidDyn* python pipeline^35^ was used to estimate the area per lipid (APL) and bilayer thickness. Both APL and thickness computing techniques allow for local definitions of these quantities, taking into account the presence of the protein and giving reliable results in the presence of membrane curvatures. The APL is obtained via a Voronoi tessellation of each lipid’s local neighborhood, while thickness is calculated using neighborhood-averaged coordinates and corresponding lipids in the opposite leaflet. Lipid positions were defined using the phosphate atom P for POPC and POPE, and the hydroxyl oxygen O3 for cholesterol.

The *MDAnalysis* package *MembraneCurvature* was used to calculate the mean curvature of the membrane leaflets.^36^ The software analyses the positions of selected atoms – typically the lipid headgroups that define the membrane surface. The positions of the reference atoms are mapped onto a discrete grid, and their heights (z-coordinates) are averaged within each grid cell. By assigning a single height to each grid point, a continuous surface can be reconstructed across the membrane. From this surface, mean and Gaussian curvature, *H* and *K*, are computed, providing complementary measures of local bending and twisting across the membrane. For each replica, the S1R–ligand complex was centered, and the equilibrated parts of each trajectory were independently analyzed to calculate the membrane’s mean curvature. The phosphate atom P of POPC and POPE and the hydroxylic oxygen O3 of cholesterol were used as the reference atoms (**Supplementary Figure 1**). 15 bins were used along the bilayer plane. The analysis was applied to the entire trajectory to yield time-averaged curvature maps.

Lipid tilt was calculated using an in-house Python script, computing the angle between the vector formed by the Z-axis of the simulation box and the atom couples O3-C25 and P-C218 for cholesterol and POPC/POPE, respectively. Lipids near protein were defined as closer than 6Å, all other lipids were considered as bulk.

### Cholesterol-protein contacts

For each frame of the trajectory, a protein residue and a cholesterol molecule were considered in contact if any atom pair between them was within 6Å. Using this per-frame contact map, for every residue and each cholesterol that ever contacted it, we identified the longest uninterrupted stretch of consecutive frames in which the contact was present; the number of frames was converted into duration using the simulation timestep. We then defined *contact persistence* as the average of longest durations across all cholesterol molecules contacting a given residue. Averaging over monomers and the three simulation replicas yielded the *average contact duration*. For structural visualization, the values of the contact duration, normalized to the longest stretch, were written into the B-factor field of a protein-only PDB file.

### Ligand-protein contacts

Protein-ligand contacts were quantified using a custom Python script built on the *MDAnalysis* library.^37,38^ A residue was classified as “in contact” with the ligand if any atom of that residue was found within 3.5 Å of any ligand atom. The contact frequency is expressed as percentage of the frames in which the contact occurs over the total number of frames in the MD trajectory.

### Correlation network analysis (CNA)

Construction and analysis of the protein dynamic network was performed with the R package Bio3D v2.4.^39^ In this approach, the protein is modelled as a network in which residues are the nodes and the correlated motions are represented as edges. The weight of the edges are calculated according to the Pearson’s cross-correlation value c_ij_, defined as:

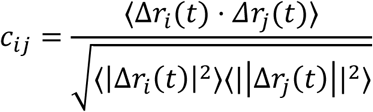

where Δ𝑟(𝑡) is the displacement of Cα of residue *i* or *j* from their average positions at time *t* and the angled brackets denotes averaging over the simulation time. The correlation of atomic displacements was calculated for each Cα pair and edges were included in the network when |𝑐_𝑖𝑗_| ≥ 0.75. To determine this cutoff, we computed the network modularity^40^ across thresholds ranging from 0.70 to 0.95. The modularity is a measure of how well a network can be divided into communities or modules, i.e., groups of nodes that are densely connected internally but sparsely connected to other groups. At lower threshold values, modularity remains stable, with residues clustering in large communities. In contrast, at higher thresholds, modularity rises steeply, leading to communities containing only a few residues – up to the limit case of one community per residue. We selected the threshold at the inflection point of this curve, where modularity began to increase steeply, in order to retain strong correlations while avoiding the formation of broad, uninformative communities. The final consensus network was constructed based on the three replicate simulations after concatenation.

Community analysis and node centrality were performed with Bio3D to characterize each network. Communities were identified by hierarchical clustering based on a betweenness clustering algorithm.^41^ The resulting communities or clusters are aggregates of nodes that are highly intraconnected but loosely interconnected. Node centrality, defined as the number of unique shortest paths crossing that node, quantifies the importance of single nodes in the constructed network.

## Results

### Ligand-bound Sigma1 receptor trimers modulate local membrane lipid organization

To investigate how membrane composition influences S1R structural dynamics, we performed all-atom MD simulations of the trimeric S1R embedded in two distinct lipid environments: a pure POPC bilayer and a MAM-mimicking membrane composed of POPC, POPE, and cholesterol (**Figures 1B** and **1C**). Each system was simulated in triplicate for 1 μs, with the receptor bound to either the agonist (+)-pentazocine or the antagonist haloperidol, yielding a total of 12 μs of trajectory data.

We assessed the structural stability of S1R during the simulations by monitoring the RMSD of the Cα atoms of the protein (**Supplementary Figure 2**). Across the three replicates, the RMSD values of S1R complexes in the MAM-mimicking membrane, after an initial equilibration period, remained stable over the course of the 1-µs simulation. On the contrary, the protein complexes in pure POPC displayed consistently higher RMSD and greater fluctuations.

To provide a molecular rationale for the observed protein stability, we analyzed global bilayer properties, including the area per lipid (APL) and bilayer thickness. These metrics enabled us to quantify the reciprocal influence between S1R and the surrounding membrane environment. Notably, the MAM-mimicking membrane showed a markedly reduced APL relative to the POPC alone (pure POPC: 49.64 ± 0.04 Å^2^ and 49.65 ± 0.04 Å^2^; MAM-like: 62.49 ± 0.02 Å^2^ and 62.70 ± 0.20Å^2^, for Hal and PnT-bound systems, respectively) (**Supplementary Figure 3A**). On the other hand, the average thickness exhibited an increase in the presence of cholesterol (pure POPC: 38.29 ± 0.02 Å and 38.21 ± 0.02 Å; MAM-like: 41.10 ± 0.04 Å and 41.19 ± 0.03 Å, for Hal and PnT-bound systems, respectively) (**Supplementary Figure 3B**). This trend was consistent across both ligand-bound systems, indicating that bilayer packing is determined primarily by membrane composition rather than ligand identity. These findings are in line with the vast literature on the condensing effect produced by cholesterol.^42,43^ The steroid nucleus restricts the conformational freedom of the lipid acyl chains and promotes their alignment, leading to reduced APL and increased membrane thickness. The condensing effect of cholesterol thus offers a structural rationale for the lower RMSD and greater stability of S1R in the MAM-like membrane.

Interestingly, both membrane types display numerous outliers characterized by low thickness values (**Supplementary Figure 3B**). Visual inspection revealed that this thinning does not arise from uniform membrane distension, which would otherwise also emerge from APL, but due to non-uniform lipid distribution in the bilayer. Specifically, regions of reduced thickness consistently form above the S1R receptor core, as the membrane deforms into a concave shape, curving inward like a pit. To quantify the topological changes in the membrane, we computed the mean and Gaussian curvature of the membrane bilayer. Curvature analysis shows positive mean curvature (H>0) and a positive Gaussian curvature (K>0) in S1R proximity, indicating the membrane curves in a bowl-shaped pit by bending in the same direction along both principal axes (**Figure 2A, 2B**; **Supplementary Figure 4**).

**Figure 2.**
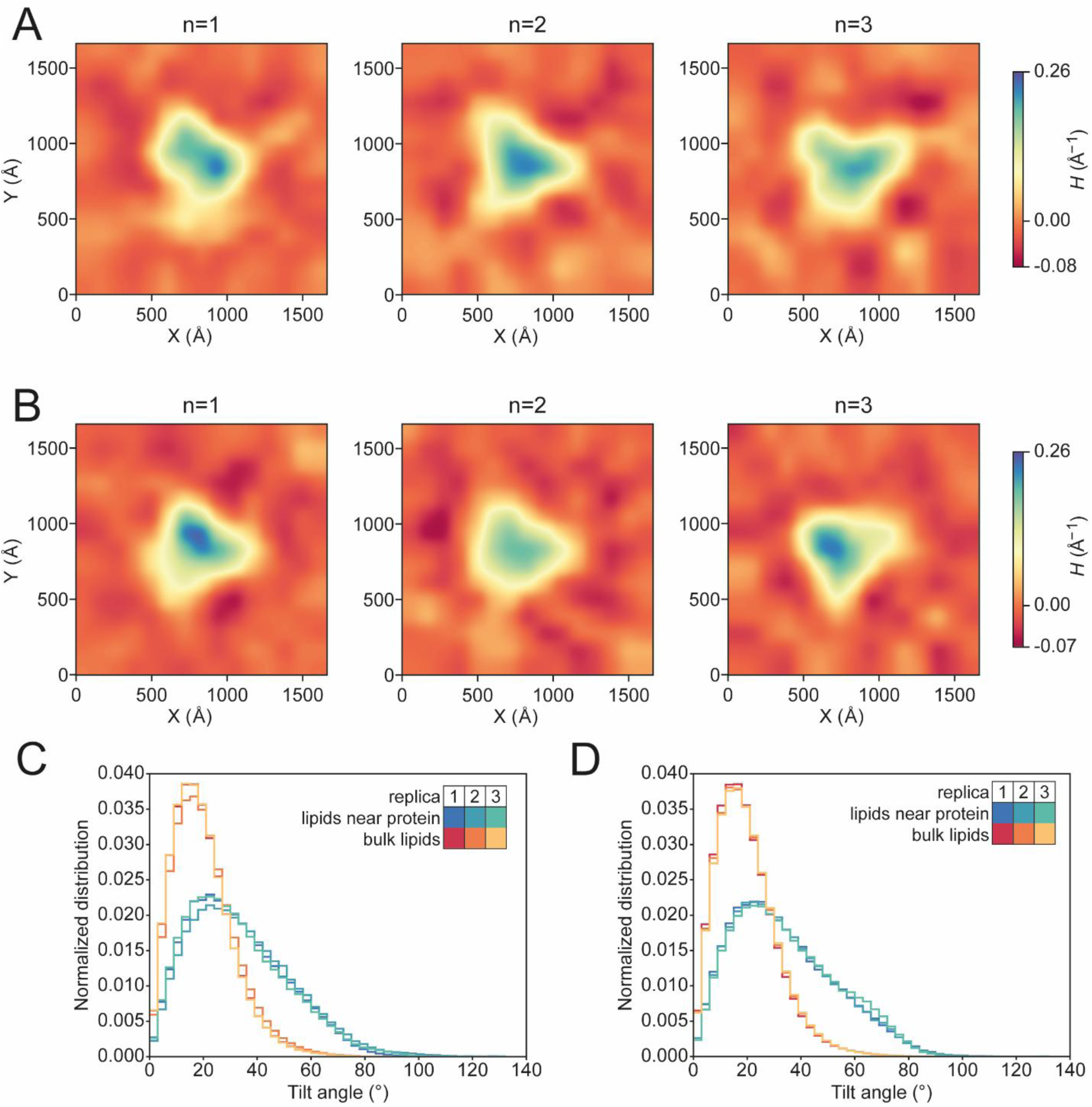
2D maps of the mean curvature (*H*) induced by S1R-PnT (**A**) and S1R-Hal (**B**) in the MAM-mimicking membrane outer leaflet in each simulation replicate (n=1, 2, 3). The curvature is calculated along the membrane surface and color coded as indicated by the scale bar. A positive curvature value indicates a concave membrane morphology with the protein occupying the pit. Distribution of the lipid tilt angles with respect to the vector normal to the bilayer surface for the S1R-PnT (**C**) and S1R-Hal (**D**) simulations. Lipids within 6Å of the protein (in blue) are more tilted compared to lipids in the bulk (in orange).

This local membrane thinning is the result of two concurrent effects: (i) an asymmetric distribution of lipids between the two leaflets, and (ii) ligand-independent spatial rearrangement of the lipids during the simulations. In the initial configuration of our system, the inner leaflet — where the receptor is embedded — contains fewer lipids than the outer leaflet (400 vs. 450, **Supplementary Table 1**). The observed asymmetry arises from the steric hindrance imposed by the hydrophobic protein platform formed by helices α4 and α5, which effectively shields the polar phospholipid headgroups from solvent exposure. This passive constraint is further reinforced by a second effect: the tilting of lipids in the inner leaflet (**Figures 2C** and **2D**). To quantify this phenomenon, we computed lipid tilt angles relative to the bilayer normal (see Materials and Methods for details). Lipids within 6 Å of the protein display a broad distribution of titles, with angles reaching up to 90°, indicating near-parallel alignment with the membrane plane. In contrast, lipids in the bulk predominantly adopt angles around 20°, consistent with a well-ordered bilayer.^44^ The enhanced tilting near S1R is a product of the direct hydrophobic interaction between the lipids acyl chains and the protein platform, which is strongly apolar in nature. Notably, the S1R receptor crystal structure already shows electron density in the area between the protein platform and the α1 helix attributable to ordered lipid heads^45^ (**Supplementary Figure 5**), supporting the propensity of this region to modulate local membrane organization as well as the robustness of our MD simulations.

### Cholesterol binds S1R in specific hotspots

Next, we focused on the lipid distribution around the S1R. It is well know that S1R interacts with cholesterol and other steroids.^46–48^ Among these, neurosteroids, synthesized within the brain and modulators of the neuronal function, have been proposed as endogenous ligands of S1R. Examples of neurosteroids are pregnonolone, dehydroepiandrosterone (DHEA) and dehydroepiandrosterone sulfate (DHEAS), that binds to S1R in the micromolar range.^46,47^ Before structural data were available, two putative steroid-binding regions were proposed based on homology to the fungal sterol C7-C8 isomerases: steroid binding domain-like I (SBDLI; residues 91-109) and SBDLII (residues 176-194).^49^ Inspection of the receptor’s crystal structure shows that SBDLII corresponds to α4 helix, while SBDLI encompasses the β2-β3 strands of cupin-like β-barrel. Together, SBDLI and SBDLII surround the protein binding site, and their involvement in the steroid recognition has been supported by the structure of *Xenopus laevis* S1R crystallized with neurosteroid ligands bound withing the β-barrel.^50^ More recently, Bezprozvanny and coworkers have identified a novel cholesterol-binding motif in the transmembrane region of human S1R, spanning amino acids 7 to 14,^15,51^ whose sequence resembles a CARC-like motif^52^ (**Supplementary Figure 5**). They demonstrated that mutations in this motif disrupt S1R clustering and alter its subcellular localization, underlining the functional importance of cholesterol interactions in regulating receptor organization.

In our MD simulations, no restraints were applied between S1R and membrane lipids, yet cholesterol molecules spontaneously accumulated in the proximity of the receptor. To identify regions of the protein that form the most stable lipid interactions, we mapped the contact persistence onto the protein structure by coloring each residue according to the average duration of its longest cholesterol-binding event (**Figure 3**). As expected, for both S1R-ligand complexes, the cholesterol interacts with parts of S1R in the membrane (α1) and those located at the protein-membrane interface (α4 and α5 helices). The most persistent contacts were observed at the CARC-like motif and at the junction between α1, α4 and α5, a region occupied by ordered lipids in the published crystal structures. Interestingly, while the CARC-like motif and the α4 helix both exhibit high contact persistence, the overall frequency of cholesterol interaction, measured as the average of the contact duration, differs between helices, suggesting distinct modes of cholesterol engagement across the protein. Cholesterol interacts most frequently with residues in α5, whereas the α1 helix shows fewer but most-long lived interactions.

**Figure 3.**
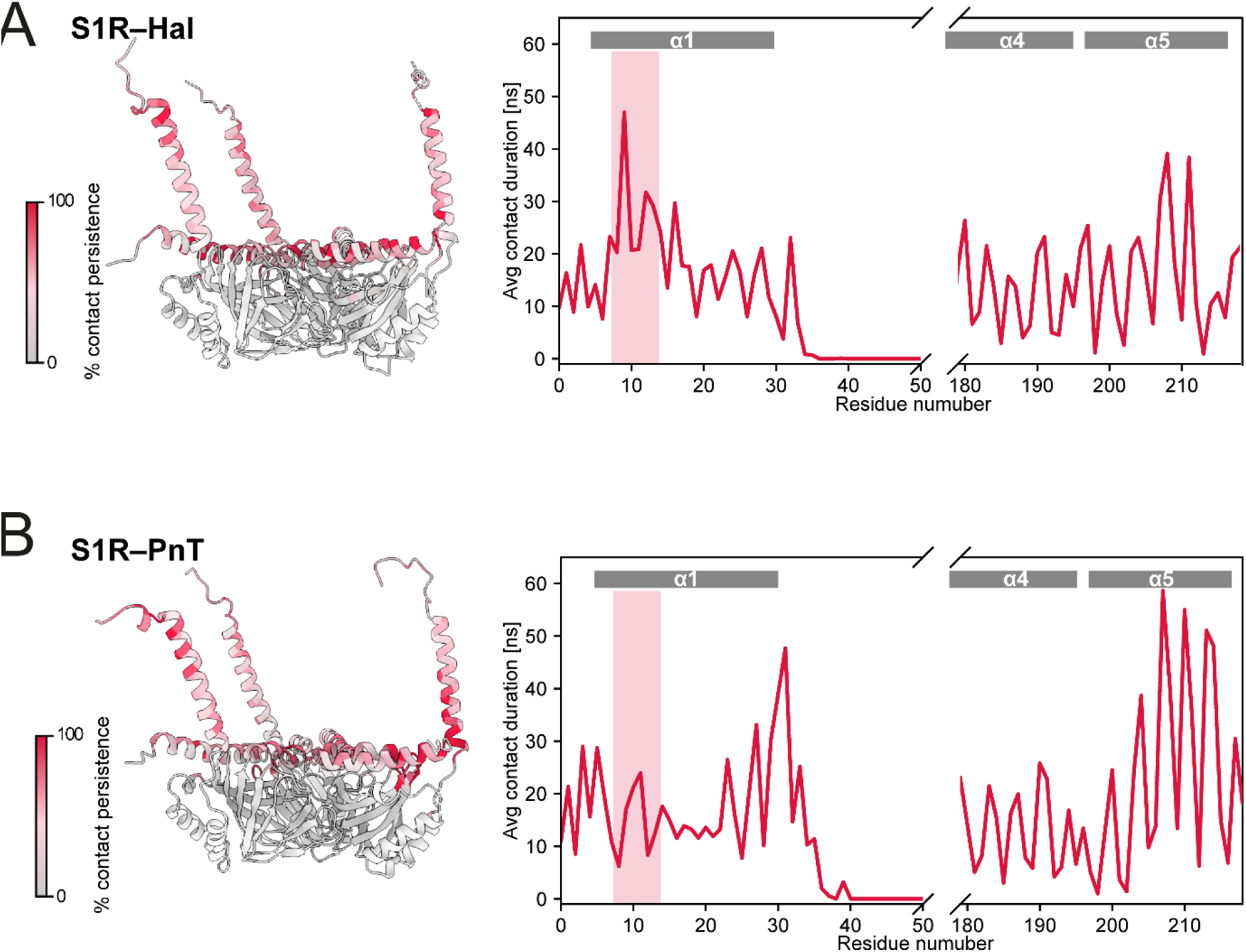
Cholesterol–S1R interaction persistence across the receptor in the S1R–Hal (A) and S1R–PnT (B) complexes. On the left-hand side, the panels show the mapping of cholesterol interaction persistence onto the S1R trimer. Each residue is colored according to the average duration of its longest cholesterol-binding event, expressed as a percentage of the maximal persistence observed in the simulation. On the right-hand side, the plot shows the protein-cholesterol contact duration averaged across the three monomers and replicas. Residue 7 to 14 belonging to the CARC-like motif are highlighted by the pink rectangle.

Taken together, these analyses demonstrate that membrane composition dictates bilayer packing, thickness, curvature, and cholesterol enrichment around S1R, whereas the nature of the bound ligand (agonist vs. antagonist) exerts negligible influence on these membrane-level properties.

### W136 emerges as a key determinant of ligand-dependent structural rearrangements

After establishing that the presence of cholesterol in the membrane plays a non-negligible role in S1R simulations, we examined the structural differences between the S1R–Hal and S1R–PnT complexes, focusing exclusively on MAM-mimicking systems, each simulated for 3 𝜇s in three replicates. Crystallographic studies have shown that the two complexes are highly similar, with an overall RMSD of 0.439Å, and both ligands engage the same key residues within the binding pocket. PnT and Hal interact with the hydrophobic side chains in the cavity and contact W89, Y103, and F107, while forming the characteristic salt bridge with E172. Schmidt and co-workers previously reported subtle differences between the complexes, including ligand-dependent effects on the helices α4 and α5.^9^

We focused on the comparison of the binding modes of the two ligands throughout the MD simulations. To this end, we quantified ligand-protein contacts throughout each trajectory (**Figure 4**). Overall, the contact profiles were highly similar, with notable differences localized mainly to residues 83-87 and 132-137. In particular, the segment spanning residues 132–137, corresponding to the β6 strand that forms the lower portion of the ligand-binding pocket, is of particular interest. This region is preferentially engaged by the chlorophenyl ring of Hal, whose more elongated structure enables deeper penetration into the cavity compared with PnT. Interestingly, the β6 strand contains W136, a residue located at the intermonomeric interface that establishes multiple hydrophobic contacts with residues of the adjacent monomer, including F83 (β1 strand), A92 (β2 strand), and T109 (β3 strand). Notably, experimental studies have shown that mutating W136 with glycine markedly impairs S1R multimerization,^5^ producing an effect closely resembling that observed upon binding with Pnt^6^. This strong agreement further supports and validates our computational findings.

**Figure 4.**
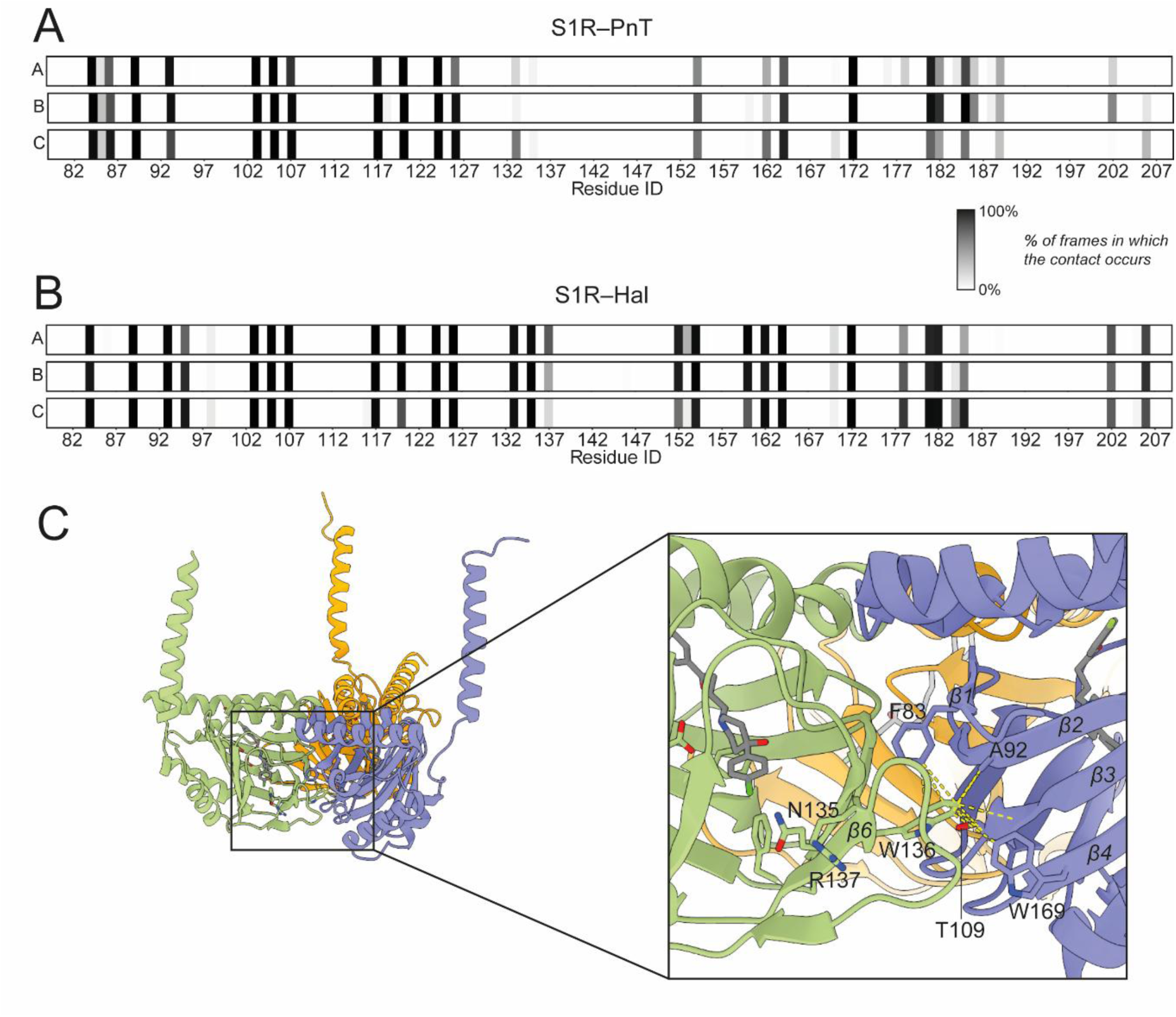
Ligand-protein contact fingerprints for the S1R-PnT and S1R-Hal complexes. Time-averaged interaction fingerprints are shown for each protomer (A, B, C) of S1R bound to (+)-pentazocine (**A**) or haloperidol (**B**). The color scale reports the percentage of simulation frames in which a ligand–residue contact is observed. Both ligands display broadly similar contact profiles, with the highest interaction frequency on E172. Notable differences emerge in the β6 region, where haloperidol forms more persistent contacts. (**C**) Zoom in on the β6 strand of the crystal structure of the S1R-Hal complex. The W136 in the β6 of one protomer forms key hydrophobic interactions with residues of the adjacent monomer (F82, A92, T109, W169).

### Ligands alter S1R interprotomer interaction and network

The different ligand-protein contact fingerprints returned by S1R-PnT and S1R-Hal complexes prompted us to analyze the MD trajectories by Correlation Network Analysis (CNA). In this framework, the protein is represented as a graph in which residues are modelled as nodes connected by edges that reflect the strength of their correlated motions. CNA therefore reveals the importance of each node in the network and allows identification of the residues with the highest degree of “centrality”, defined as the number of edges associated to each node.

Across both complexes, residue W136 emerges as the major hub of the network (**Figure 5A**). However, its centrality is markedly higher in S1R-Hal and the overall centrality distributions differ in the two systems. In S1R-PnT, a broader set of residues displays high centrality values, reflecting a more fragmented and distributed community architecture. Consistently, community detection reveals 24 distinct communities in S1R-PnT, compared with only 14 in S1R-Hal. This difference indicates that the Hal-bound receptor features more cohesive and tightly interconnected communication pathways, whereas the PnT-bound system exhibits a less integrated signaling network (**Figure 5B**). We calculated the shortest paths connecting two selected nodes; the paths provide information on how the dynamic processes are propagated through the protein. We selected as starting point of the paths (the so-called source) E172, a key interaction point for all S1R ligands, while residue W169 of the adjacent monomer was selected as end point (the sink). It has been reported that tryptophan residues, such as W169, play a unique role in the protein-protein interaction thanks to their ability to contribute to aromatic interactions, to act as hydrogen bond donors and to their large hydrophobic surface.^53–55^ To compare the overall coupling strength between the source and the sink, we calculated the path length distribution for the S1R–Hal and S1R–PnT. The length is defined as the sum of the edge weights along the shortest communicative route between two residues in protein network. Shortest paths, such as those observed for S1R–Hal, are associated with a strong coupling (**Figure 5C**). Structural rearrangements are transmitted through the protein backbone of the β10-strand (from E172 to R175) and via the β-barrel loop (Y120, W121, A159, T160) are focalized on W136. From there, all of the 100 paths that we computed cross residues T109, A110 connecting W136 to W169 of the adjacent monomer.

**Figure 5.**
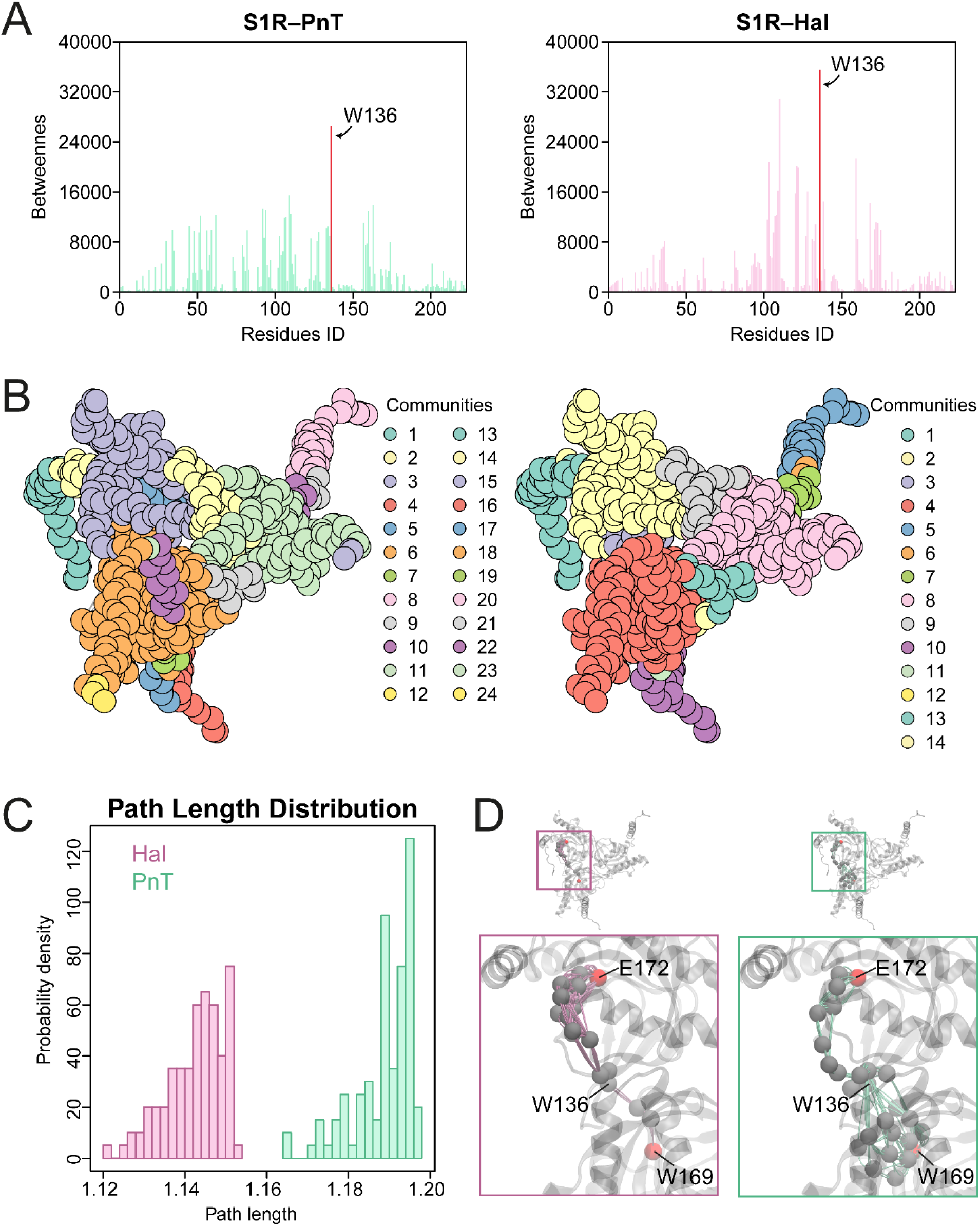
CNA analysis of the MD simulations of S1R–PnT and S1R–Hal. (A) Network centrality of the Cα atoms in S1R–PnT and S1R–Hal. (B) Nodes of S1R–PnT (left) and S1R–Hal (right) are depicted as circles colored according community they belong to. (C) Probability density distribution of the path lengths at the intermonomeric interface (with source/sink pair E172 and W169) (D) Representation of 100 communication pathways are viewed as lines on the protein structure (S1R–PnT, green; S1R–Hal, pink). The “source” and “sink” residues are colored in red.

In S1R–PnT, the path behavior is radically different. Paths cross the β-barrel over the hydrogen bonds connecting the β-strands (from E172 on β10-strand, to S125 on β5-strand) to T160 on β8-β19 loop in a straightforward manner. Then, when the paths cross the gap between one monomer and the adjacent one, they split incoherently in different directions.

## Conclusions

In this study, we combined all-atom MD simulations with network-based analyses to investigate how membrane composition and ligand identity shape the structural dynamics of S1R. We found that the physical properties of the membrane – particularly the presence of cholesterol – plays an important role in modulating S1R structural stability. Cholesterol condenses and thickens the bilayer, promotes more ordered membrane packing, and contributes to the maintenance of a stable trimeric assembly, independently of the ligand bound to the receptor.

In contrast, the agonist (+)-pentazocine and the antagonist haloperidol produce localized and mechanistically distinct effects within the receptor core. While both ligands interact with the canonical binding pocket, only haloperidol engages the β6 strand more deeply and consistently, stabilizing W136 — a residue located at the interprotomer interface. This stabilization strengthens intermonomeric coupling and reorganizes the receptor’s internal communication network into a more cohesive and efficient architecture. Conversely, (+)-pentazocine weakens β6-mediated coupling, resulting in a more fragmented interaction network and more heterogeneous communication pathways across the trimer, consistent with the experimentally observed agonist-driven destabilization of higher-order oligomers.

Together, these findings provide actionable guidance for ligand design at the S1R receptor. By revealing that agonists and antagonists differentially modulate the β6 strand and its key residue W136, our analysis identifies a structural switch that medicinal chemists can directly target. Elongated molecules interacting with the β6 strand should stabilize interprotomer coupling. These insights connect ligand scaffolds to predictable oligomeric outcomes, offering a practical framework for designing molecules with tailored functional profiles.

## ASSOCIATED CONTENT

### Data Availability Statement

The raw data associated with this work are freely available on a Zenodo repository at the following link: https://doi.org/10.5281/zenodo.18787104.

### Supporting Information

Figure S1: Structure of cholesterol (CHOL), phosphatidylcholine (POPC) and phosphatidylethanolamine (POPE) – the lipids that compose the membrane in which the S1R receptor is embedded. Relevant atom nomenclature is reported. The black dashed line represents the vector used to calculate the lipid tilt angles.

Figure S2: Root-mean square deviation (RMSD) of the protein Cα over the course of the simulations. RMSD was computed relative to the first frame after alignment to the entire protein. Panels A shows the S1R-PnT and S1R-Hal in POPC, whereas panel B reports the corresponding systems in POPC/POPE/CHOL.

Figure S3: Boxplot with quartiles, median and outliers of the lipid-averaged area per lipid (APL) (A) and bilayer thickness (B), computed across all replicates for the four simulated systems. The POPC membrane (in red and light blue) exhibits a higher APL and a correspondingly lower bilayer thickness compared with the POPC/POPE/CHOL system (in pink and light green).

Figure S4: Analysis of the membrane curvature induced by S1R–PnT and S1R–Hal in the MAM-mimicking (A) and in POPC (B) membrane outer leaflet, in each simulation replicate (n=1, 2, 3). Top row: 2D maps of the lipids’ head (P atom) z-coordinate variation from lowest z-coordinate value (*Z*). Middle row: 2D maps of the membrane mean curvature (*H*). Bottom column: 2D maps of the membrane gaussian curvature (*K*). (C) Geometric interpretation of the curvature values *H* and *K*.

Figure S5: S1R receptor modulates local membrane organization. (A) Ordered lipids are observed in the crystal structures 6dk1 (left) and 6djz (right) at the junction of the luminal core and the α1 helix (shown in yellow sticks, oxygen atoms in red). Protein ligands (+)-pentazocine and haloperidol are shown in blue and pink, respectively. (B) Steroid-binding regions of S1R are highlighted on the S1R-progesterone structure (8w4b): two steroid binding domain-like regions (SBDLI and SBLII) and CARC-like motif.

Table S1: Membrane composition for S1R-Hal and S1R-PnT systems

Table S2: Simulation box details

Table S3: Gromacs simulation parameters employed for trajectories production

## ACKNOWLEDGMENTS

This work was funded by the National Research Council (CNR–Italy) under the program “PROGETTI DI RICERCA @CNR” (acronym DATIAMO), by the European Union (Next Generation EU) under the program PRIN of the Ministero Dell’Università e della Ricerca (Progetti di ricerca di Rilevante Interesse Nazionale 2022 – CODICE 2022Z3BBPE – CUP: B53D23015780006 – Development of broad-spectrum coronavirus antiviral agents acting as allosteric modulators of the host protein sigma-1) and by the project “Potentiating the Italian Capacity for Structural Biology Services in Instruct Eric”, acronym “ITACA.SB” (Project No. IR0000009, CUP B53C22001790006), funded by the European Union’s NextGenerationEU under the MUR call 3264/2021 PNRR M4/C2/L3.1.1. We acknowledge ISCRA for awarding this project access to the LEONARDO supercomputer, owned by the EuroHPC Joint Undertaking, hosted by CINECA (Italy).

## Notes

### Competing Interest Statement

The authors have declared no competing interest.

