## Supplementary Information for "MD simulations of Human Sigma1 Receptor Trimer Uncovers Cholesterol Dependent Stabilization and Ligand Specific Dynamics"

**Supplementary Tables**

**Table S1:** Membrane composition for S1R-Hal and S1R-PnT systems

**Table S2**: Simulation box details

**Table S3**: Gromacs simulation parameters employed for trajectories production

**Figure S1**

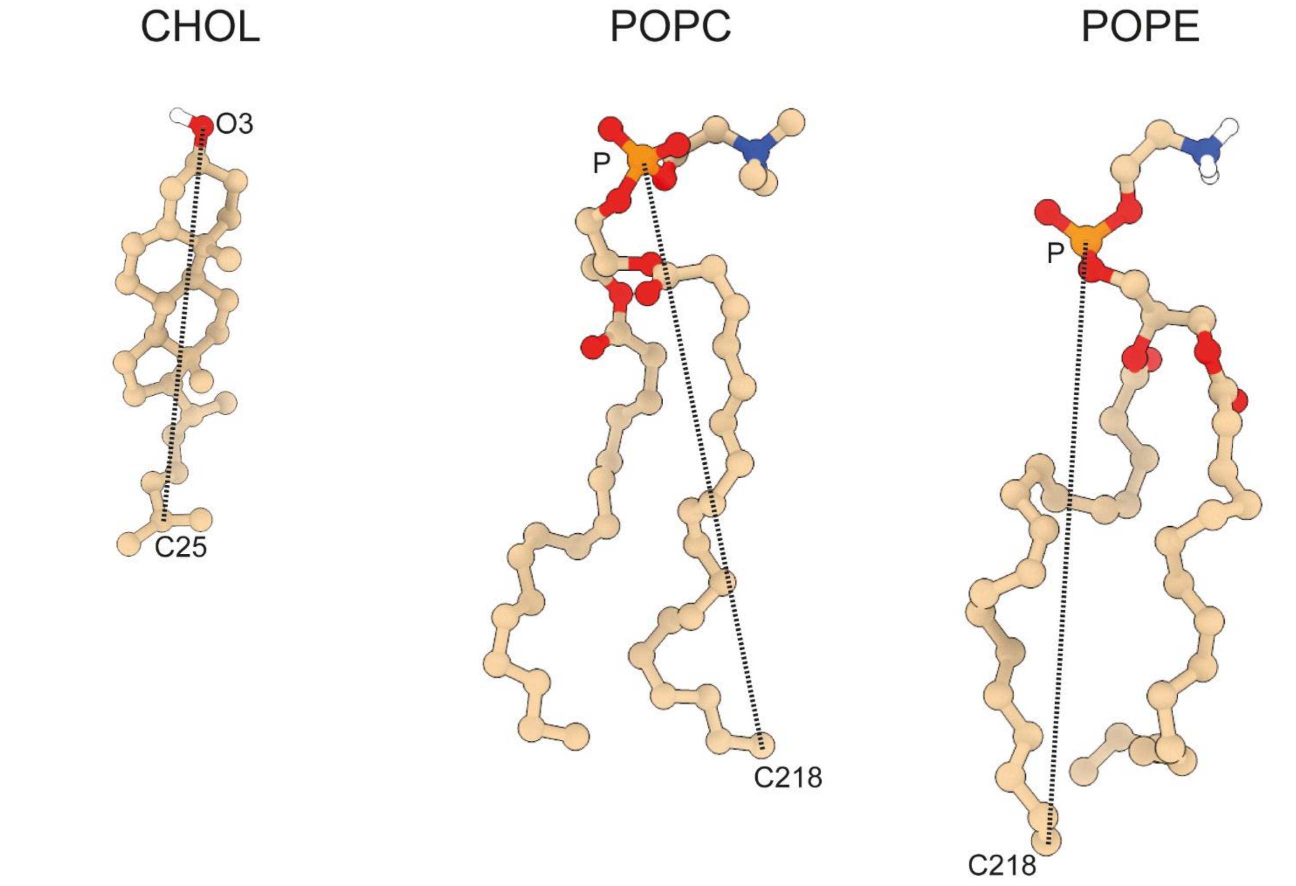

**Figure S2.**

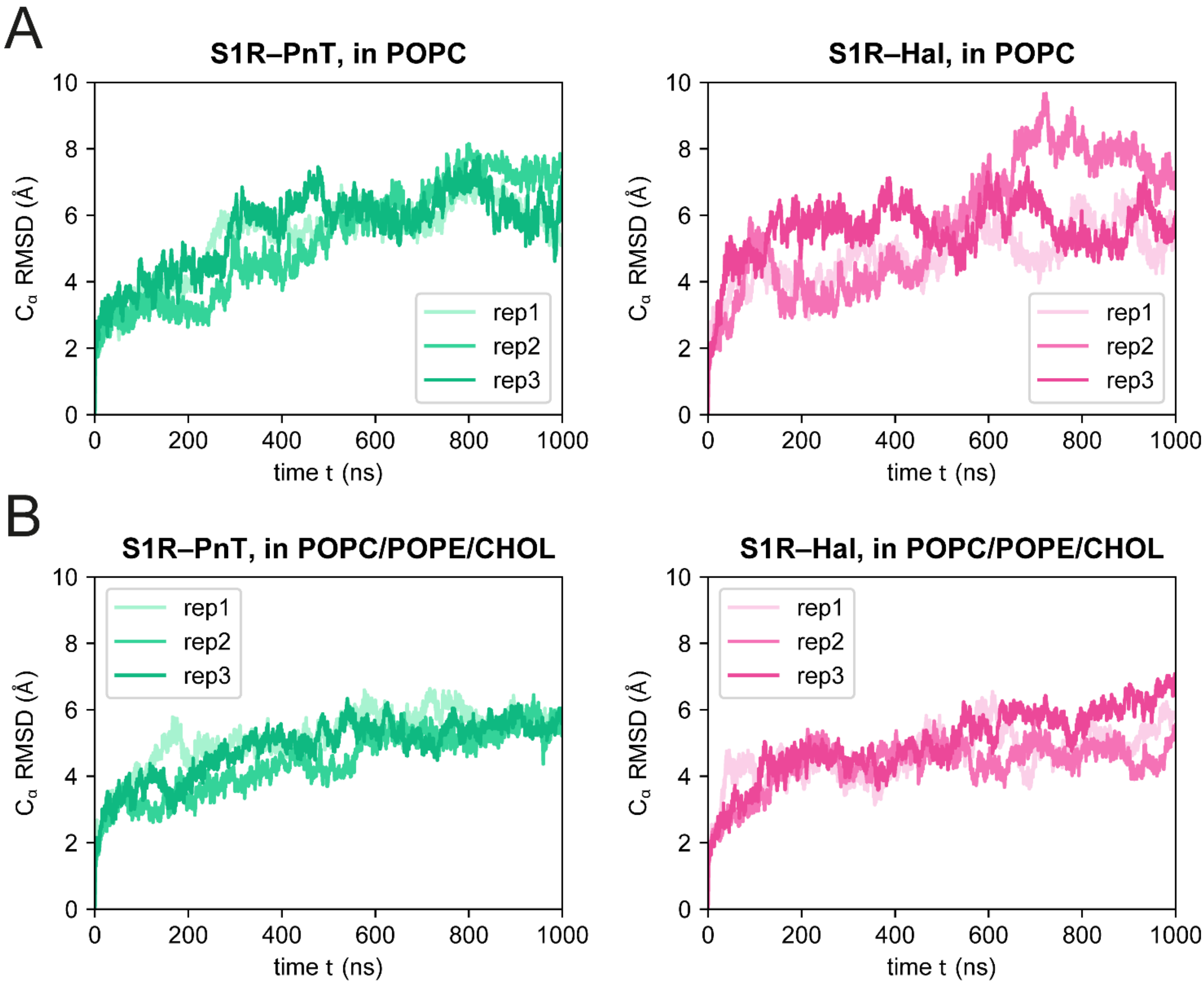
A
S1R–PnT, in POPC
S1R–Hal, in POPC
B
S1R–PnT, in POPC/POPE/CHOL
S1R–Hal, in POPC/POPE/CHOL
Cα RMSD (Å)
time t (ns)
rep1
rep2
rep3
0
200
400
600
800
1000
0
2
4
6
8
10

**Figure S3.**

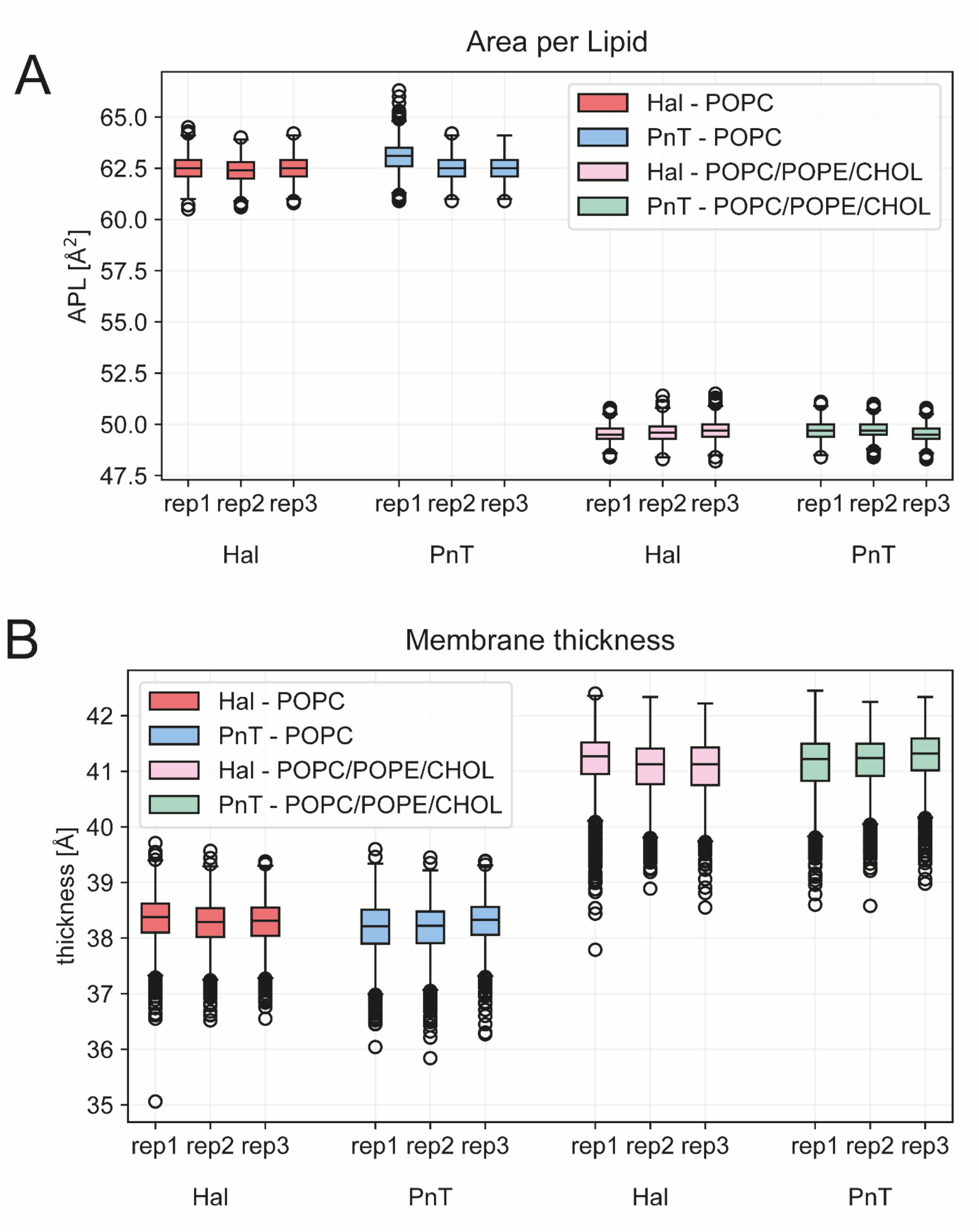

A
Area per Lipid
APL [Å²]
65.0
62.5
60.0
57.5
55.0
52.5
50.0
47.5
Hal - POPC
PnT - POPC
Hal - POPC/POPE/CHOL
PnT - POPC/POPE/CHOL
rep1 rep2 rep3
rep1 rep2 rep3
rep1 rep2 rep3
rep1 rep2 rep3
Hal
PnT
Hal
PnT
B
Membrane thickness
thickness [Å]
42
41
40
39
38
37
36
35
Hal - POPC
PnT - POPC
Hal - POPC/POPE/CHOL
PnT - POPC/POPE/CHOL
rep1 rep2 rep3
rep1 rep2 rep3
rep1 rep2 rep3
rep1 rep2 rep3
Hal
PnT
Hal
PnT

**Figure S4.**

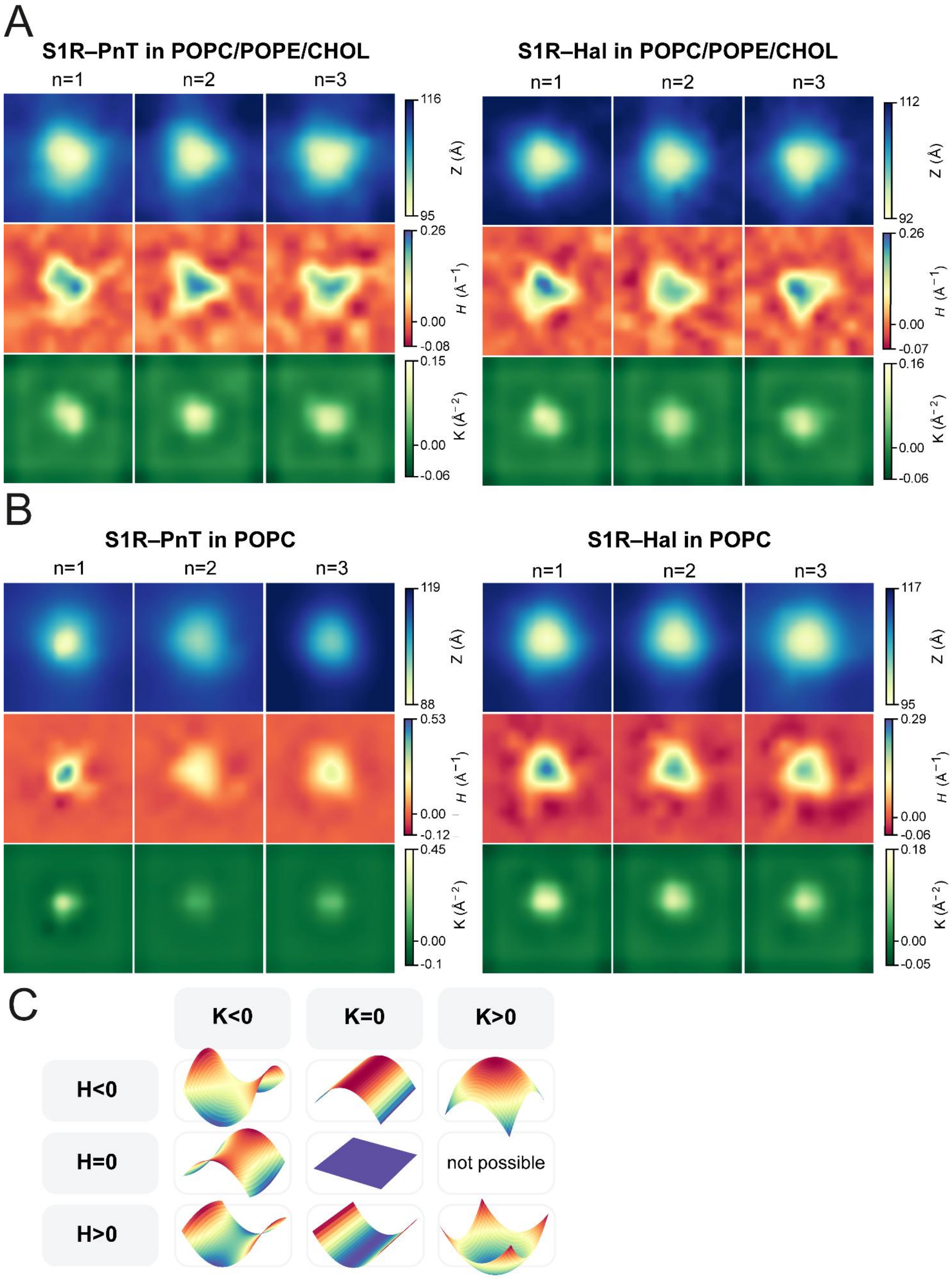
A
S1R–PnT in POPC/POPE/CHOL
S1R–Hal in POPC/POPE/CHOL
n=1
n=2
n=3
116
95
Z (Å)
0.26
0.00
-0.08
H (Å⁻¹)
0.15
0.00
-0.06
K (Å⁻²)
112
92
0.26
0.00
-0.07
0.16
0.00
-0.06
B
S1R–PnT in POPC
S1R–Hal in POPC
119
88
0.53
0.00
-0.12
0.45
0.00
-0.1
117
95
0.29
0.00
-0.06
0.18
0.00
-0.05
C
K<0
K=0
K>0
H<0
H=0
H>0
not possible

**Figure S5.**

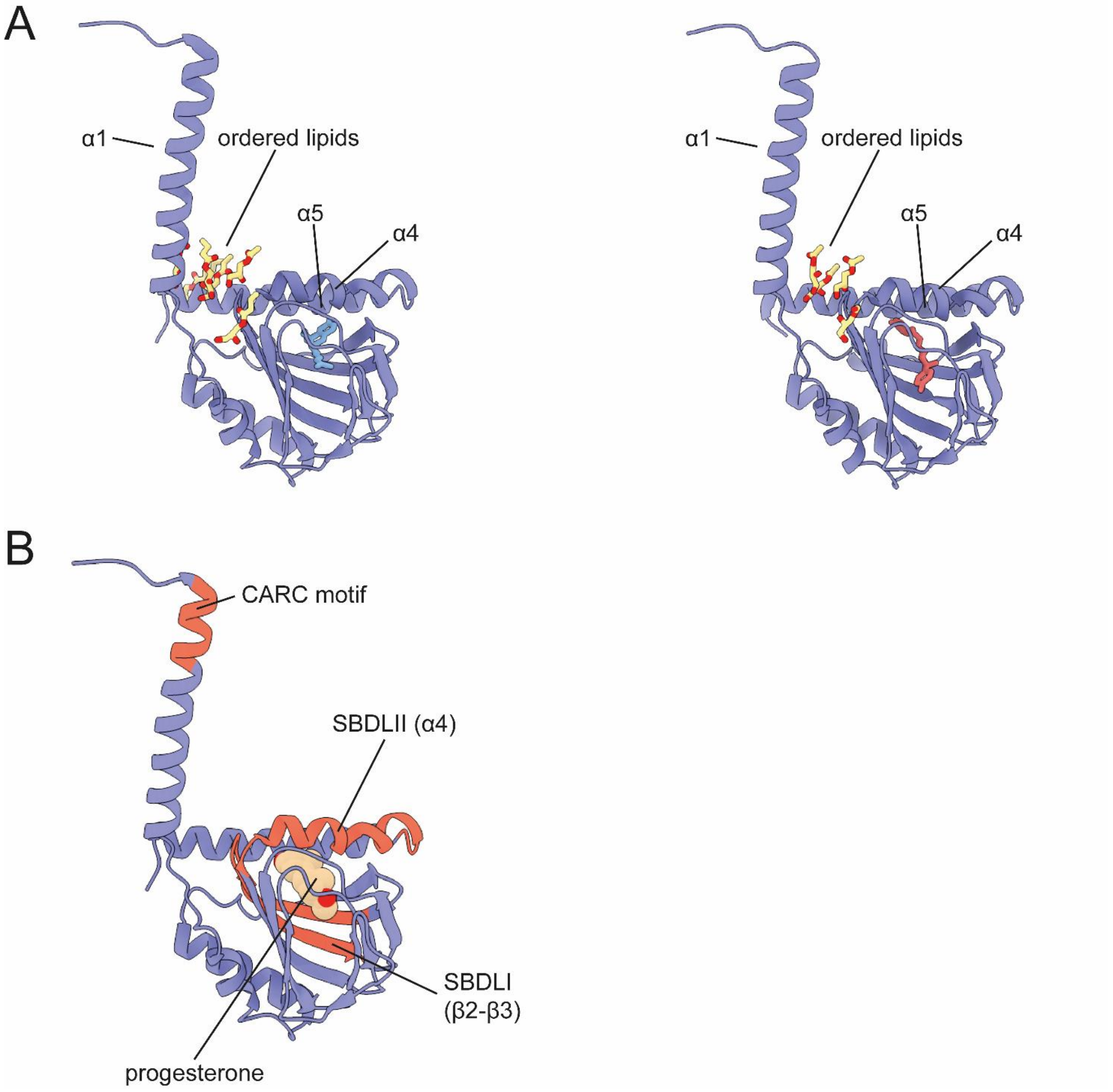
A
α1
ordered lipids
α5
α4
α1
ordered lipids
α5
α4
B
CARC motif
SBDLII (α4)
SBDLI
(β2-β3)
progesterone

**Table 1.**

| System | Leaflet | POPC | CHOL | POPE | TOT LIPIDS |
|---|---|---|---|---|---|
| S1R-Hal in POPC | Inner | 326 | 0 | 0 | 326 |
| | Outer | 365 | 0 | 0 | 365 |
| S1R-PnT in POPC | Inner | 325 | 0 | 0 | 325 |
| | Outer | 364 | 0 | 0 | 364 |
| S1R-Hal in MAM-like membrane | Inner | 224 | 88 | 88 | 400 |
| | Outer | 252 | 99 | 99 | 450 |
| S1R-PnT in MAM-like membrane | Inner | 224 | 88 | 88 | 400 |
| | Outer | 252 | 99 | 99 | 450 |

**Table 2.**

| | S1R-Hal in POPC | S1R-PnT in POPC | S1R-Hal in MAM-like membrane | S1R-Hal in MAM-like membrane |
|---|---|---|---|---|
| Water mol | 78884 | 81341 | 85084 | 88011 |
| Lipid mol | 691 | 689 | 850 | 850 |
| $Na^+$ mol | 227 | 234 | 246 | 255 |
| $Cl^-$ mol | 217 | 224 | 236 | 245 |
| Tot atoms | 340507 | 347618 | 367548 | 376341 |
| Initial box dimensions (x, y, z) [nm] | 16.0,16.0,14.3 | 16.0,16.0,14.6 | 16.6,16.6,14.3 | 16.6,16.6,14.6 |

**Table 3.**

| Parameter | Value |
|---|---|
| integrator | md |
| dt | 0.002 |
| nstxtcout | 25000 |
| nstcalcenergy | 100 |
| nstenergy | 1000 |
| nstlog | 1000 |
| cutoff-scheme | Verlet |
| verlet-buffer-tolerance | -1 |
| nstlist | 50 |
| rlist | 1.33 |
| vdwtype | Cut-off |
| vdw-modifier | Force-switch |
| rvdw_switch | 1.0 |
| rvdw | 1.2 |
| coulombtype | PME |
| rcoulomb | 1.2 |
| pbc | xyz |
| tcoupl | v-rescale |
| tc_grps | SOLU MEMB SOLV |
| tau_t | 1.0 1.0 1.0 |
| ref_t | 310 310 310 |
| nsttcouple | 50 |
| pcoupl | C-rescale |
| pcoupltype | semiisotropic |
| tau_p | 5.0 |
| compressibility | 4.5e-5 4.5e-5 |
| ref_p | 1.0 1.0 |
| nstpcouple | 50 |
| constraints | h-bonds |
| constraint_algorithm | LINCS |
| continuation | yes |
| nstcomm | 100 |
| comm_mode | linear |
| comm_grps | SOLU_MEMB SOLV |
